# Cultural evolution by capital accumulation

**DOI:** 10.1101/707620

**Authors:** Jean-Baptiste André, Nicolas Baumard

## Abstract

In this article, we model cultural knowledge as a capital in which individuals invest at a cost. To this end, following other models of cultural evolution, we explicitly consider the investments made by individuals in culture as life history decisions. Our aim is to understand what then determines the dynamics of cultural accumulation. We show that culture can accumulate provided it improves the efficiency of people’s lives in such a way as to increase their productivity or, said differently, provided the knowledge created by previous generations improves the ability of subsequent generations to invest in new knowledge. Our central message is that this positive feedback allowing cultural accumulation can occur for many different reasons. It can occur if cultural knowledge increases people’s productivity, including in domains that have no connection with knowledge, because it frees up time that people can then spend learning and/or innovating. We also show that it can occur if cultural knowledge, and thus the higher level of resources that results from increased productivity, leads individuals to modify their life history decisions through phenotypic plasticity. Finally, we show that it can occur if technical knowledge reduces the effective cost of its own acquisition via division of labour. These results suggest that culture should not be defined only as a set of knowledge and skills but, more generally, as all the capital that has been produced by previous generations and that continues to affect current generations.

---

In economic terms, the soma of biological organisms constitutes a *capital*, an asset that enhances the organism’s ability to perform useful activities (Kaplan et al., 2000, 2003). Organisms invest in it because it allows them to better exploit the environment and ultimately to reproduce more. Biological growth is therefore a form of capital accumulation. But biological capital is often limited in its lifespan: it is *disposable* in the sense that it is not passed onto the next generation (Kirkwood, 1977). In many species and at least in all mammals, natural selection has favored a life history strategy whereby organisms build their biological capital and then let it degrade (Jones et al., 2014). Only a very small fraction of an organism’s biological capital, its germ cells, survives the death of the organism and is passed on to its offspring. For all the rest, each generation must start from scratch.

But this is not the case for all forms of capital built by biological organisms. Some forms of capital are functionally independent of the biological body, and can therefore retain value after the death of the individual who produced them. That is, they are *heritable*. For these investments, each generation can start a little higher than the former.

Investments in knowledge are the most typical forms of heritable investments. New knowledge, not always but in many cases, remains useful long after the death of the person who produced it. But knowledge is not the only form of heritable capital. All the investments made in improving the quality of the environment (sometimes called niche construction; Laland et al., 2000) can be heritable. Clearing part of a forest, building a house, opening a path, fighting infectious diseases, etc., are all heritable investments. In this article we focus primarily on knowledge, but to understand its dynamics, we argue that it must be seen as one form, among others, of investments in heritable capital.

Like investments in biological growth, investments in the production and social acquisition of cultural capital have a cost for individuals (Boyd and Richerson, 1985, Lumsden and Wilson, 1981, Rogers, 1988). First, in the vast majority of cases, innovations are the result of efforts made by individuals to solve a problem (Allen, 2009, Bloom et al., 2017, Mokyr, 1992; and see Fogarty et al., 2015 for a discussion). Even the inventions that famously have a strong serendipitous component (e.g. the discovery of penicillin, microwave ovens, velcro, saccharin, superglue, etc.), are always the work of engineers, researchers, or more generally of people who pay attention to an accidental observation, and put enough effort to transform it into an actual discovery. Second, acquiring knowledge socially is not costless either. During their growth, humans invest in the development of their behavioral skills at the expense of other investments, resulting in a delay of reproduction (Kaplan et al., 2000). In Drosophila, Mery and Kawecki (2004) experimentally showed that learning has an operating cost: Drosophila placed in conditions where they must learn have a reduced egg laying rate.

There is a rich series of models in the literature that explicitly consider the costs of innovation and social learning, and study the evolution of individuals’ investment strategies and their consequences on the dynamics of cumulative culture (Aoki et al., 2012, Kobayashi et al., 2019, Lehmann et al., 2013, Mullon and Lehmann, 2017, Ohtsuki et al., 2017, Wakano and Miura, 2014). However, this line of models undertake a specific assumption regarding social learning: they assume that the speed of social learning increases linearly with the amount of knowledge available that an individual does not yet not possess. Even though this assumption may sometimes be appropriate, it has no reason to be true in general, and it happens to have important consequences regarding the dynamics of cultural accumulation. In a highly original approach, Mesoudi (2011) shows that, without this assumption, the cost of social learning constitutes a constraint that limits the total amount of knowledge that a population is able to maintain. In any case, nevertheless, these models do not consider the competition between investments in heritable versus non-heritable –that is, disposable– capital.

In this article we consider cultural knowledge as a heritable capital voluntarily produced by individuals by paying a cost, and we take into account the fact that individuals can also invest in non-heritable capital. The rate of cultural evolution thus eventually depends on the fraction of their investments that individuals make in heritable versus non-heritable capital. To characterize these investment decisions, we assume that individuals have been shaped by natural selection to invest at all times in the form of capital that has the strongest marginal effect on their fitness. This approach, typical of the study of life history strategies (see e.g. Lehmann et al., 2013), allows us to capture the essential mechanisms that affect individuals’ investment decisions, and ultimately to describe the dynamics of cultural evolution.

In the cultural evolution literature, culture can accumulate because it allows individuals to acquire knowledge and skills more efficiently than if they had had to create them from scratch (e.g. Aoki et al., 2012, Boyd and Richerson, 1995, Enquist et al., 2007, Enquist and Ghirlanda, 2007, Lehmann et al., 2013). That is, pre-existing knowledge is a facilitator. As a result, it gives the individuals of subsequent generations a head start and ultimately allows them to go further than the individuals who preceded them. The first and central outcome of our article is to show that this principle must in fact be generalized. Cultural accumulation can occur provided culture facilitates the life of individuals, and this can happen for many reasons, of which reducing the cost of acquiring knowledge is just one example. Culture can accumulate if it increases the amount of resources individuals invest in knowledge, the efficiency of their investments into innovation, or even if it merely facilitates their investments in non-cultural activities. In all these cases, the knowledge available to a given generation eventually improves the ability of individuals to invest in new knowledge, allowing them to go further than the previous generation.

Second, in addition to making cumulative culture possible in a broader set of circumstances, we find that this generalized view of culture as capital also results in a broader range of possible cultural dynamics. Depending on the precise effects of knowledge on individuals’ vital rates, cultural accumulation may be infinite or finite, it may experience one or more phases of acceleration, and culture may even have several equilibria. Hence, our model helps explain well-established observations on the historical dynamics of knowledge (Aiyar et al., 2008, D’Errico and Stringer, 2011, Lehman, 1946, Mokyr, 1990, and see also Enquist et al., 2008).

Third, our results also shed light on the origins of variations in the speed of cultural evolution (see also Henrich, 2004, Kolodny et al., 2015, 2016, Vaesen et al., 2016). On one hand, we show that demography has little effect on the speed and extent of cultural accumulation. Adding more individuals to a population, all unable to innovate beyond existing knowledge, does not foster the accumulation of more knowledge. This helps understand, for instance, why the industrial revolution took place in relatively small but rich countries, rather than in the most densely populated but poorer ones (Baumard, 2018, Clark, 2008, McCloskey, 2016). On the other hand, in our model, the respective rates of cultural evolution in different fields of knowledge all depend similarly on the time and resources that individuals have at their disposal to innovate. As a result, all these rates move in parallel, including in intellectually unrelated fields. Our results thus help understand why independent areas of knowledge often flourish, stagnate, or regress together in the same historical periods.

To understand our results intuitively, it is useful to think of Dino Buzzati’s short story *The Eiffel Tower* (Buzzati, 1966). Buzzati imagines that, when his tower has reached the 325 metres planned by the government, Gustave Eiffel proposes to his workers to raise it higher, without limits. Workers accept and the tower grows taller and taller, but a practical problem eventually slows down its rise. To add metal beams to the top of the tower, every morning, workers have to walk up the hundreds of metres already built, and every night they have to walk down them again. Eventually, there comes a day when the tower is so high that, in their working day, workers have just enough time to climb it up and down and no time left to actually work. The tower cannot grow any higher.

Like Buzzati’s workers over the course of a day, each person over the course of their life must learn, or at least take advantage, of all the cultural knowledge already available, and only then can they contribute to increasing this knowledge further. When the amount of knowledge becomes too high, the cost of acquiring it becomes so large that people simply do not have the time and resources available to create more. Like Buzzati’s tower, cultural knowledge reaches a ceiling.

Things would be different, however, if the tower was not only a tower but a tool that would enable workers to be more efficient. If, for instance, workers also built an elevator, then they would have invested into a capital that would reduce (rather than increase) the cost of climbing up to the top. To accumulate significantly, knowledge must increase, not reduce the possibility of producing more knowledge, by reducing the cost of reaching the technological frontier and/or by increasing people’s ability to innovate. In this case, then not only can cultural knowledge accumulate, but its accumulation can also accelerate, rather than slow down.

## Models and Results

### Overview of the modelling approach

We build a microscopic model of cultural accumulation. Our objective is to describe the dynamics of the amount of knowledge –and more generally the dynamics of heritable capital– as a macroscopic consequence of investments made by individuals during their growth. To simplify the analysis, we represent the amount of heritable capital present in a population as a one-dimensional quantity, *z*. This approach is inspired by life history models in which all the dimensions of individual growth are summarized into a single variable measuring the amount of biological capital of each individual (often called their body weight). This unidimensional simplification is also undertaken by several scholars in the field of cultural evolution (e.g. Enquist et al., 2008, Lehmann et al., 2013) as well as in endogenous growth models in economics (Romer, 1986).

The basic building block of our models consists in the description of the growth of individuals, from birth to death, as a succession of investments (see also Lehmann et al., 2013). During this growth, individuals invest in biological capital, by growing their soma. They also invest in capital outside their body, by improving their environment. And they invest in knowledge, both by acquiring knowledge already produced by others and also, potentially, by producing new knowledge themselves. We distinguish these various forms of capital according to a single property that we assume, for the sake of simplicity, binary: capital can be heritable or non-heritable. The first consists in all the investments that retain a value after the biological death of the individual, while the second consists in the investments whose value disappears upon the death of the individual.

At each age *a* of her life, an individual is characterized by the quantities *x*(*a*) and *y*(*a*) of non-heritable and heritable capital that she owns, respectively, at this age. With this capital, the individual produces a certain amount of resources that she can then decide to allocate to three possible activities.(1) First, she may invest in the growth of her non-heritable capital, *x*(*a*). We will call this type of investment “growth.” (2) Second, she can invest in the production of new forms of heritable capital, *y*(*a*), by producing new knowledge, or by improving the quality of her environment. We will call this type of investment “innovation” (even if it really covers a broader set of activities). Note that, in the cultural evolution literature, this type of investment is usually called individual learning. Here we prefer using the term innovation to highlight the fact that new knowledge is produced rather than simply learnt. (3) Third, she can invest in the growth of her heritable capital, *y*(*a*), not by producing new forms of heritable capital by herself, but by converting the heritable capital already produced by others into her own heritable capital, typically by learning existing knowledge. We will call this third type of investment “social learning” or most often simply “learning” (even if, here too, it potentially covers a broader set of investments). We define three parameters that determine the efficiency of converting resources in these three activities respectively. The efficiency of growth is *γ*, meaning that, per unit of resources invested in growth, *x* increases by *γ*, while λ and *η* measure the efficiency of social learning and innovation, respectively. Throughout the article we make the standard assumption that learning socially a certain amount of knowledge is always more efficient than producing this same knowledge from scratch, i.e. that *η* < λ (see e.g., Lehmann et al., 2013, Mullon and Lehmann, 2017, Ohtsuki et al., 2017, Wakano and Miura, 2014), which entails that individuals always start by learning all available knowledge first –because this is more efficient– and innovate only in a second step, as in critical social learning (Enquist et al., 2007).

Throughout our analysis, in the sake of simplicity, we make the assumption that individuals have an exogenously determined time budget *L* for their development (that is, for investing in growth, learning, and/or innovation), after which they stop developing and start investing exclusively in reproduction. In other words, this entails that we assume (1) that individuals follow a bang-bang strategy whereby they first allocate all their resources to the development of their capital, and then switch to an exclusive investment in reproduction and (2) that the switching age (that is, the age at sexual maturity) is exogenously constrained and does not evolve in our model. As a consequence of this simplifying assumption, we do not consider here the trade-off between growth and reproduction (unlike Lehmann et al., 2013 or Wakano and Miura, 2014 for example). Instead, we consider the trade-off between investing in knowledge (social learning or innovation) and investing in the growth of non-cultural capital.

At each instant of their life, individuals are characterized by a production function *P* (*x, y*) measuring the rate at which they produce resources (that can then be invested into various activities), in function of their amounts of heritable and non-heritable capital, *y* and *x*. Throughout this article, as is often the case in economics, we will use the Cobb-Douglas production function

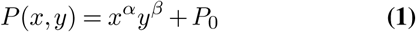

where the parameters *α* and *β*, both strictly positive, represent so-called elasticities of production, that is the responsiveness of production to a change in either *x* or *y*. We will always consider the case where *α* + *β* ≤ 1, which means that total capital has diminishing returns (or at most a constant return). We will often use the variable 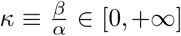, that represents the relative importance of heritable capital in an individual’s production function. That is, *κ* represents the relative importance of heritable capital (e.g. knowledge) among the set of needs of individuals. The parameter *P*_0_ represents essentially the resources obtained from parental care. In the course of the analyses we will often use the partial derivatives of this production function. Hence, we define 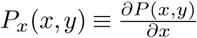 and 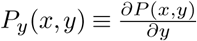.

We chose a Cobb-Douglas function because it provides an incentive for individuals to always invest a non-zero fraction of their resources in each type of capital, and because this fraction naturally depends both on the efficiency of each capital and on the efficiency of the investments to increase this capital. On the other hand, as any hypothesis, the choice of this particular function introduces some limitations to our analysis. In particular, the Cobb-Douglas function has infinite partial derivatives in zero for each type of capital, which cannot be right biologically speaking for a production function. However, it should be noted that, in this model, the Cobb-Douglas function determines the growth rate of individuals, and not a demographic growth rate. No individual reproduces at an infinite rate in our model.

We assume that individuals’ allocation decisions have been shaped by natural selection so that they make optimal allocation decisions at each age. This hypothesis cannot be strictly valid in all circumstances, but it is useful at least as a first approximation. Standard result of optimal control theory shows that the optimal allocation consists in making, at all times, the investment(s) with the greatest marginal effect on the production function (Iwasa and Roughgarden, 1984, Perrin, 1992, Perrin et al., 1993). This allows us to describe the series of investments made by an individual over the course of her life (see also Lehmann et al., 2013). Our model is deterministic with discrete generations. We consider a Wright-Fisher population of constant size, in which all individuals are born at the same time, and grow in parallel.

Let us consider a population at the beginning of generation *t*, with an amount of cultural knowledge (or more generally an amount of heritable capital) equal to *z*_*t*_. Individuals of this generation develop by making optimal investment decisions over a period of time *L*. Among other things, they may learn socially the existing knowledge *z*_*t*_ (from the adults of the previous generation), and/or they may produce new knowledge on their own. At the end of this development, individuals reach a given level of non-heritable capital, *x*(*L*), and a given level of knowledge, *y*(*L*), which thus becomes the knowledge available for the next generation, that is *z*_*t*+1_ = *y*(*L*). The objective of our model is to describe the dynamics of *z*_*t*_ from generation to generation.

### Model 1 - The burden of knowledge and the limits of cultural accumulation

The objective of our first model is to characterize the dynamics of cultural evolution under the null assumption that culture does not affect the life history of individuals. To do so, we assume that the actual amount of resources produced by an individual per unit of time –that she can invest in the three possible activities, growth, learning, and innovation– is a constant independent of her capital *x* and *y*. Without loss of generality, we assume that this constant productivity is equal to 1. In this model, therefore, the production function (eq. 1) does not affect the growth of individuals. It only plays the role of a target that individuals seek to maximize at all times, which naturally determines their optimal ratio of investments in growth, learning, and innovation.

Consider a population of *N* individuals born with an initial level of *x* = 0 and *y* = 0, in a social environment in which they have a pre-existing amount of knowledge *z*. Under the assumption that social learning is more efficient than innovation, the optimal allocation schedule consists in the three following phases (see Fig. 1, and Supporting Information (SI) 1 for more details).

**Fig. 1.**
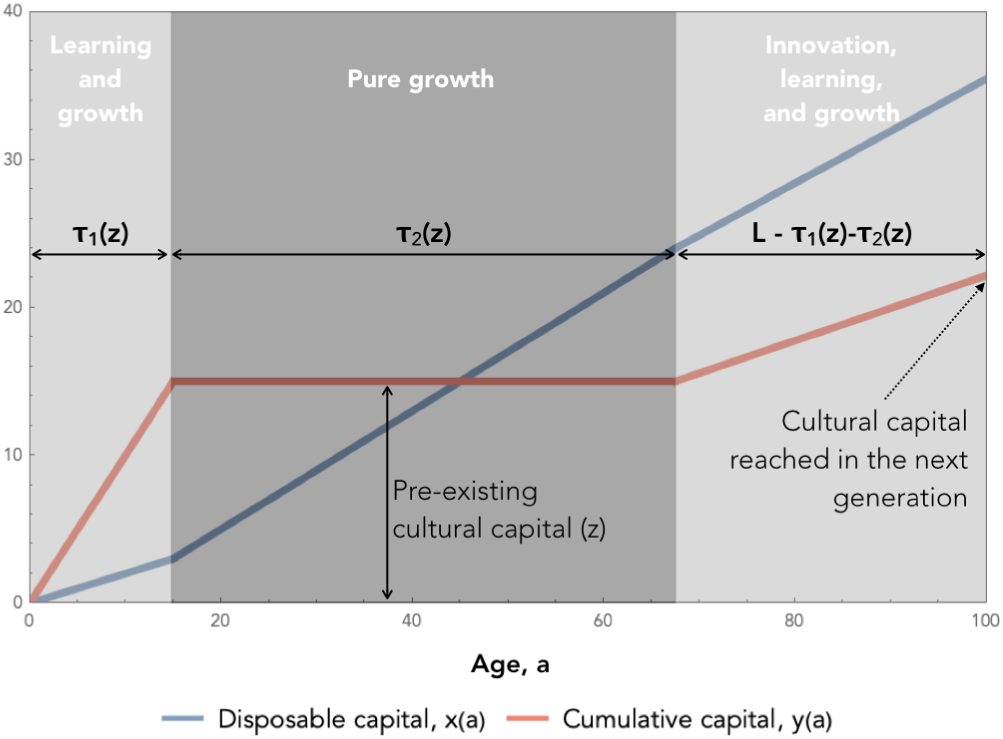
Individual development in model 1. Variations of the two types of capital through the development of an individual when the amount of pre-existing heritable capital is *z*. Here, parameters are chosen to make the figure as readable as possible: *γ* = 0.4, *λ* = 2, *η* = 0.25, *α* = 0.5, *β* = 0.5, *N* = 100, *L* = 100, *z* = 15

#### Learning and growth toward the frontier

During the first phase of their lives, individuals invest in parallel in the growth of their personal capital *x* and the learning of the available knowledge *z* (i.e. in the conversion of public capital *z* into personal capital *y*). In SI 1.1.1, we show that this phase lasts for a time 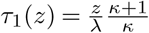

#### Pure growth toward the frontier

Once *z* is fully socially learned, the capital *y* can potentially continue to increase if individuals invest in innovation. However, since the efficiency of innovation is lower than the efficiency of social learning this implies that, at the end of Phase 1, investing in growth is necessarily more profitable than innovating. So individuals do not start innovating right away. They first experience a phase during which they only invest in the growth of *x*. They do so until the value of investing in *x* becomes as low as the value of innovating, at which point they start innovating. In SI 1.1.2, we show that this second phase lasts for a time 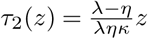. Hence, catching-up takes a total time *τ* (*z*) ≡ *τ*_1_(*z*) + *τ*_2_(*z*), which allows us to define the average growth rate of individuals during the catch-up phase

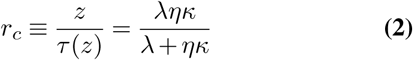

#### Innovation at the frontier and cultural interference

Once the frontier is reached, individuals start investing in innovation. But they are not alone. Other individuals in the population also invest in innovation in parallel. Here we assume that individuals are aware of the innovations made, or in the process of being made, by others. As a result, since it is more efficient to learn something socially than to reinvent it by oneself, individuals always decide to focus on innovations that are not already made by others, which results in a sharing of the innovation work. This does not mean that we assume individuals to be cooperative, however. On the contrary, individuals always behave so as to maximize their own self-interest, but they do so by taking into account others’ behavior. In SI 1.1.3, we show that the rate at which a population of *N* individuals produces new knowledge under this assumption is

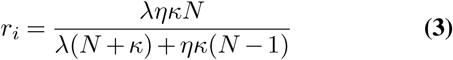

This shows that population size, *N*, has a positive but saturating effect on the rate of cultural evolution at the frontier (the derivative of *r*_*i*_ with respect to *N* is positive but decreasing, and it tends towards 0 when *N* tends towards infinity). As a result, the rate of cultural evolution increases with *N* but asymptotically tends towards a maximum 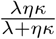 for large population sizes. This is a consequence of the fact that individuals must spend time learning the innovations made by others, which forces them to divert resources away from their own innovations. In other words, if more heads means more ideas, it also implies more ideas to learn, and therefore less time available to create new ideas. In parallel with the notion of selective interference, which limits the speed of biological adaptation, we call this phenomenon “cultural interference.”

**Table 1.** List of symbols used in main text.

| Symbol | Definition |
| --- | --- |
| $a$ | Age of an individual |
| $x(a)$ | Non-heritable capital of an individual of age $a$ |
| $y(a)$ | Heritable capital of an individual of age $a$ |
| $N$ | Population size |
| $z_t$ | Heritable capital available in a population at the end of generation $t$ |
| $\alpha$ | Exponent of $x$ in the Cobb-Douglas production function |
| $\beta$ | Exponent of $y$ in the Cobb-Douglas production function |
| $\kappa$ | $\beta/\alpha$ |
| $P_0$ | Production independent of personal capital |
| $P(x, y)$ | Production of an individual with capital $x$ and $y$ |
| $P_x(x, y)$ | Marginal productivity of $x$ for an individual with capital $x$ and $y$ |
| $P_y(x, y)$ | Marginal productivity of $y$ for an individual with capital $x$ and $y$ |
| $l(a)$ | Fraction of resources invested in social learning at age $a$ |
| $d(a)$ | Fraction of resources invested in innovation at age $a$ |
| $\gamma$ | Increase of $x$ per unit of resource invested in growth |
| $\lambda$ | Increase of $y$ per unit of resource invested in social learning |
| $\eta$ | Increase of $y$ per unit of resource invested in innovation |
| $L$ | Total duration of development |
| $\tau_1(z)$ | Duration of the first development phase when the available knowledge is $z$ |
| $\tau_2(z)$ | Duration of the second development phase when the available knowledge is $z$ |
| $\tau(z)$ | $\tau_1(z) + \tau_2(z)$ |

#### The limits of cultural accumulation

If a population contains an amount of knowledge *z* in a generation, in the next generation it contains an amount

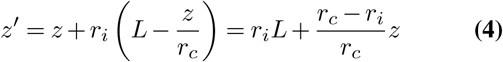

where *r*_*c*_ and *r*_*i*_ are given respectively by equations 2 and 3 above, resulting in a linear recurrence equation that can be solved and plotted (see eq. 14 in SI, and Fig. 2). This shows three things.

**Fig. 2.**
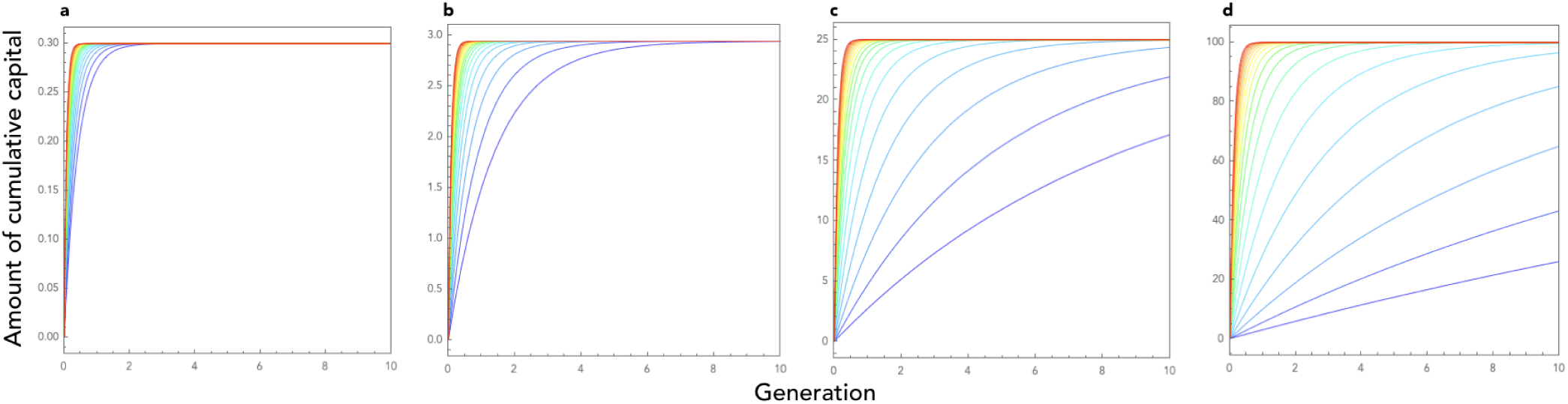
Cultural dynamics in model 1. Here we plot the dynamics of heritable capital through generations for 20 different values of population size ranging from *N* = 1 (blue curve) to *N* = 2200 (red curve). Note that the curves are drawn as if cultural evolution was a continuous phenomenon even though, in reality, generations are discrete in our model. Parameters are as follows: *γ* = 1, λ = 5, *η* = 0.1, *P*0 = 0.1, *L* = 30, and *κ* = 0.1 (**a**), *κ* = 1 (**b**), *κ* = 10 (**c**), and *κ* = 100 (**d**).

First, the level of knowledge contained in a population converges towards an equilibrium, a cultural carrying capacity, independent of population size and given by

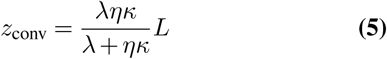

In addition to the total time budget available, *L*, this equilibrium depends on two limiting factors: the learning efficiency, λ, and the product *ηκ*, which depends both on the efficiency of investments into innovation, *η*, and on the relative importance of knowledge in the needs of individuals, *κ*. Even if the cost of social learning is not limiting (for example because λ is very large), the equilibrium amount of culture in a population is still limited by the fact that people waste time investing in non-heritable capital. Every investment in (i) learning existing knowledge and (ii) growing non-heritable capital constitutes a loss for culture, as it must be re-made from scratch every generation. It must also be noted that this equilibrium depends on the efficiency of innovation, *η*, only provided *κ* is not infinite. In fact, *η* plays a role in the equilibrium level of culture by changing the relative attractiveness of investing in innovation as compared to investing in *x*. Therefore, if one ignores the existence of non-heritable capital (i.e. if *κ* is infinite) this effect disappears.

Second, cultural accumulation is always decelerating (see Fig. 2) since the amount of newly produced culture from one generation to the next 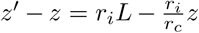 is a linearly decreasing function of *z*. This is a consequence of the fact that the technological frontier always recedes with the amount of existing knowledge, and this for two reasons. First, existing knowledge must be learnt before individuals start innovating, which takes time. Second, the more knowledge there already exists, the less further innovations are useful as compared to non-heritable capital and, therefore, the more individuals grow their *x* before they start innovating. Third, this allows us to understand the conditions that make the existence of cumulative culture possible. Cultural accumulation occurs if the amount of cultural knowledge attained in a population in a given generation increases with the amount of cultural knowledge available in the previous generation. That is, if *z*′ is an increasing function of *z*. From equation 4, we can see that this is the case provided that *r*_*i*_ < *r*_*c*_. In other words, culture gradually accumulates provided the growth rate of individuals during catch-up is larger than their growth rate during the innovation phase. More precisely, equation 4 shows that the parameter that governs the cultural accumulation potential in a population is the ratio 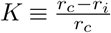, which is given by

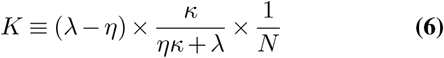

This cumulative potential depends on three factors (see also Fig. 2):

- First, *K* increases with the difference between the efficiency of learning and the efficiency of innovation, λ − *η*. This is the most obvious factor. Culture accumulates on the condition that the cost of learning what has already been discovered is lower than the cost of discovering them initially.
- Second, *K* increases with the proportion of knowledge in individuals’ capital, *κ*. The higher the knowledge content of individuals’ capital, the greater the cumulative potential. This second effect can also be understood for the same reason. When *κ* is low, the majority of the capital of individuals has to be reconstructed anyway at each generation from scratch, so there is no strong advantage given by the pre-existence of knowledge and culture is then little cumulative.
- Lastly, *K* decreases with population size *N*. Culture accumulates provided population size is not too large. This counter-intuitive result can be understood by observing that, when *N* ≫ max (1, *κ*), the rate of cultural evolution during the innovation phase is approximately equal to the rate of evolution during the catch-up phase (*r*_*i*_ ≈ *r*_*c*_; eqs. 2 and 3). In a large population, even if innovation is less efficient than social learning (that is, even if λ > *η*), the vast majority of knowledge acquired by an individual is acquired through social learning, and not through innovation, anyway. Each individual needs to produce only a very small part of the knowledge by himself. It is therefore almost the same for an individual to grow up in a population in which there is already a quantity of information *z* produced by previous generations (and where the rate of cultural growth is therefore *r*_*c*_), as to grow up in a large population that produces this information on the fly by innovation (and where the rate of cultural growth is therefore *r*_*i*_ ≈ *r*_*c*_). Culture can therefore not accumulate in an infinite population. In contrast, in a small population, innovation is a limiting factor and therefore the rate of cultural evolution is significantly slower at the frontier than during catch-up, which leads to a gradual accumulation of knowledge.

### Model 2 - Culture accumulates because it is a capital

The above model actually allows to go further. Culture can accumulate, as we said, provided that *z*′ is an increasing function of *z*. From the equation 4, we see that this can in fact be the case owing to two, apparently distinct, mechanisms.

The first mechanism, the only one considered in model 1, is the fact that, under some conditions, we may have *r*_*c*_ > *r*_*i*_. That is, social learning is sometimes more efficient than innovation, which is the central notion regarded as the heart of cumulative culture in the literature (e.g. Aoki et al., 2012, Boyd and Richerson, 1995, Enquist et al., 2007, Enquist and Ghirlanda, 2007, Lehmann et al., 2013). Here, this mechanism allowed us to define the cumulative potential of a population in model 1 (eq. 6). But equation 4 allows us to see that another mechanism can also lead to cultural accumulation, namely the fact that, under some conditions, the amount of available culture, *z*, might have positive effects on the very parameters *L, r*_*c*_, and/or *r*_*i*_ of the equation. Using the expressions for *r*_*c*_ and *r*_*i*_ (eqs. 2 and 3), we see that this second mechanism takes place if culture positively affects the amount of time that individuals devote to their growth, *L*, the efficiency of their social learning, λ, the efficiency of their investments in innovation, *η*, and/or the fraction of their capital that they invest in culture, *κ*. In other words, culture can accumulate if it is not neutral, that is if it modifies the vital rates of individuals in such a way that it increases their cultural productivity.

In reality, however, these two mechanisms are not distinct. Rather, the first is a special case of the second. According to this mechanism, the culture of a population benefits individuals by allowing them to acquire skills and knowledge more rapidly, and at a lower cost, than if they had had to invent them themselves. As a result, culture can accumulate because it allows individuals to go further than they would have had time to do if they had started from scratch. Therefore, even in model 1, culture is actually a capital that makes life easier for individuals. It just happens that this capital acts in a specific way, by reducing the cost of acquiring knowledge. Yet, there is no reason why the only way for culture to accumulate is this one. More generally, culture can accumulate because it improves the efficiency of people’s lives in many different ways, of which reducing the cost of acquiring knowledge is just one example. The reason why culture accumulates is, therefore, more general than what is usually considered in the literature. This generalization is of great importance because it allows to account for a wider range of phenomena in cultural evolution.

First, the special-case mechanism only works in small populations where innovation is actually a limiting factor, whereas the generalized mechanism can operate in principle at all population sizes, including in infinite populations. Second, while cultural accumulation under the special-case mechanism is necessarily decelerating due to the ever-increasing burden of knowledge (Fig. 2), under the generalized mechanism, cultural evolution needs not always decelerate, since the beneficial effects of the cultural capital on the vital rates of individuals can, in principle, always offset the burden of knowledge, so that the amount of newly produced culture from one generation to the next can remain constant, or even increase with *z*. The generalized mechanism can therefore lead to a broad range of cultural dynamics. Depending on the precise effects of knowledge on individuals’ vital rates, the accumulation may be infinite or finite, it may experience one or more phases of acceleration, and culture may even have several equilibria.

In the following, we aim to illustrate how these generalized effects of culture could occur. To this aim, we build a second model, similar to the first, but in which the productivity of individuals depends on their capital. As in the first model, individuals develop by accumulating two types of capital, *x* and *y*, and they are characterized at each moment of their lives by a production function given by equation 1. But here, unlike in the first model, this production is invested in the growth of *x* and *y*. If an individual of age *a* has amounts of capital *x*(*a*) and *y*(*a*), respectively, and invests a fraction *d*(*a*) of his production in innovation, a fraction *l*(*a*) in learning, and a fraction 1 − *d*(*a*) − *l*(*a*) in growth, *x*(*a*) and *y*(*a*) experience instantaneous increases:

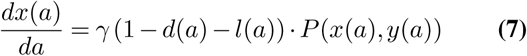

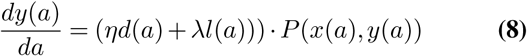

Like in model 1, at the end of their growth, individuals have reached an amount of knowledge, *y*(*L*), which then becomes the knowledge *z*, available for social learning by the next generation, and so on.

#### Optimal growth schedule

As in model 1, we assume that individuals have a time *L* to develop (after which they reproduce) and we assume that they allocate their production optimally at all times to growth, learning, and innovation, which implies that they always make the investment(s) that have the greatest marginal effect on production (see SI 1.2). This optimal allocation consists of three phases, as in model 1. The only difference with model 1 is that, here, the growth of individuals is not linear. This growth cannot be expressed analytically, hence we simulate it numerically (see SI 1.2). As in model 1, but numerically here, we derive the level of knowledge reached by a population of *N* individuals at generation *t* + 1 as a function of the level of knowledge at generation *t*.

Unlike in the first model, knowledge –or more generally heritable capital– can accumulate here from generation to generation even in a very large population (see Figs. 3 and 4). This occurs because knowledge allows individuals to be more efficient in the production of new knowledge. In result, knowledge first accumulates in an accelerating way. However, under this simple version of the model, the time it takes to reach the frontier nevertheless keeps on increasing as knowledge accumulates. At some point therefore, individuals do not have enough time left to innovate. In result, the production of new knowledge eventually decelerates, until knowledge finally reaches an equilibrium.

**Fig. 3.**
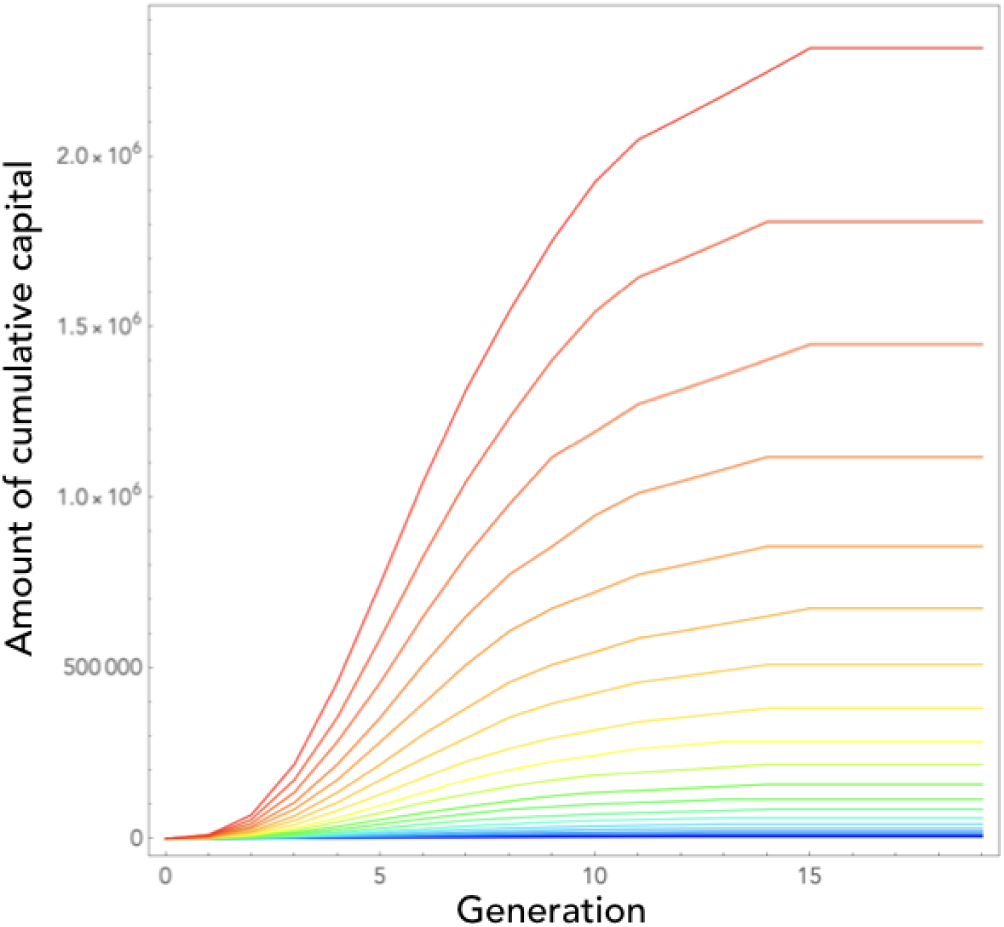
Cultural dynamics in model 2. Here we plot the dynamics of heritable capital through generations for 20 different values of population size ranging from *N* = 1 (blue curve) to *N* = 2200 (red curve). Parameters are as in figure 2, with the time-step used in numerical simulations *δ* = 0.01

**Fig. 4.**
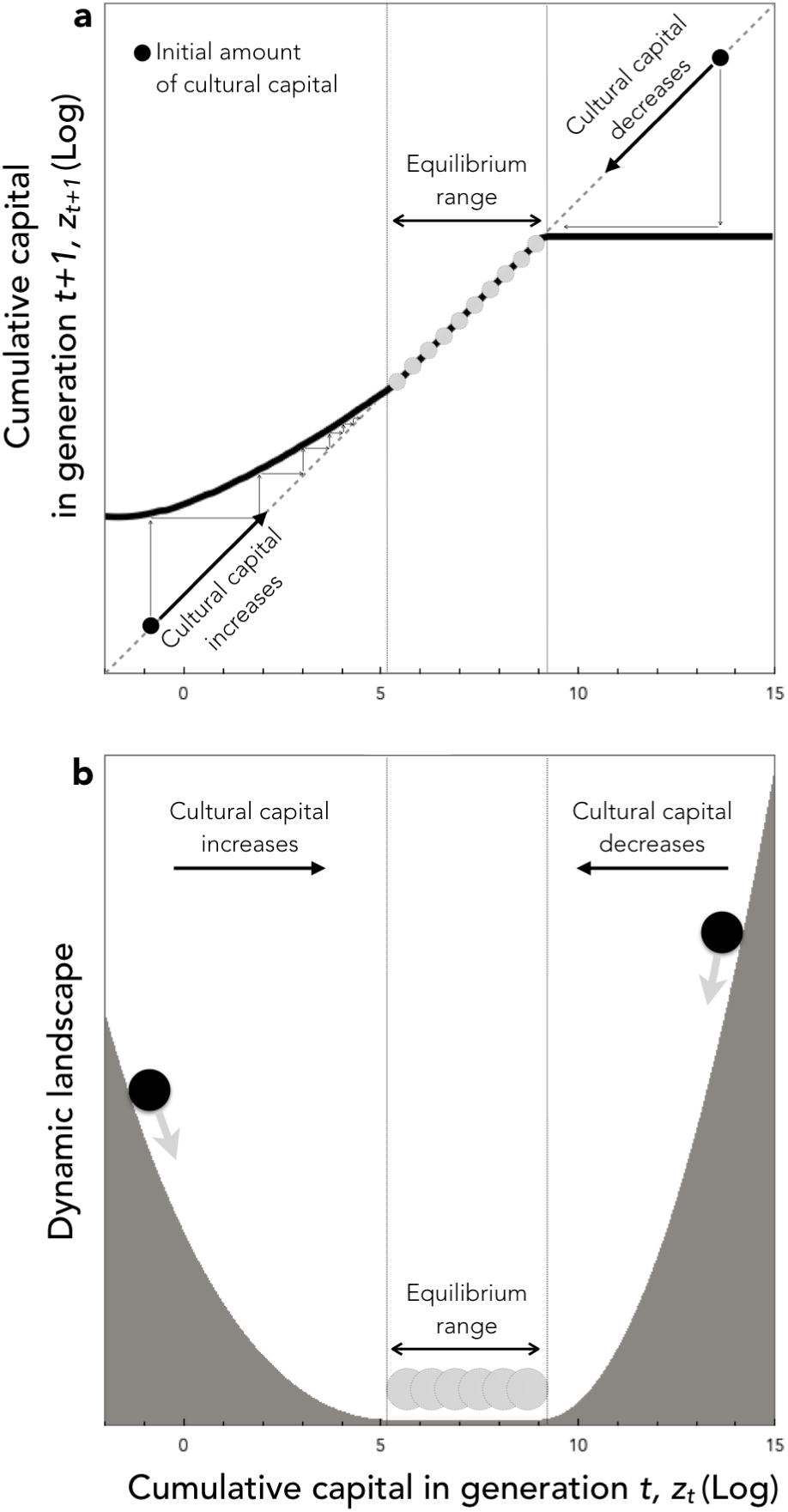
Cultural evolution in model 2. (a) The thick solid line shows the amount of cultural capital in generation *t* + 1 plotted in function of cultural capital in generation *t*. The dashed line shows the identity function. This allows to graphically represent the dynamics of culture from any initial condition, by directly reading, at each generation, the amount of culture reached in the next generation (see the thin arrows). When culture is initially low, it increases. When culture is initially high, it decreases. In between, culture has a range of intermediate equilibria. (b) Graphic representation of cultural dynamics through an analogy with a physical landscape. This allows an intuitive perception of the dynamics by picturing the course of a ball going down the hills and stopping in the plain. The landscape is constructed by setting its slope for each value of *zt* as being proportional to the change of cultural capital in one generation when starting from *z_t_*. Parameters are as in figure 3, with population size *N* = 1000

This model also allows us to explore more subtle effects of culture on individuals’ life histories. In the following, we explore three biologically plausible effects and show that they often lead to more complex cultural dynamics.

#### Investments in knowledge increase along the pyramid of needs

It is reasonable to assume that the most basic needs of individuals are purely personal and not heritable. Individuals at the beginning of their growth invest mainly in their biological capital, that will be lost upon their death. It is only when they become richer, moving upward on their so-called pyramid of needs (Maslow, 1943), that they can begin to invest in forms of capital that are further away from biology, improving their environment at a larger scale, or investing in knowledge (see Baumard, 2018 for a review). Here, we capture this effect by assuming that the ratio *κ* is low when the amount of non-heritable capital, *x*, owned by an individual is below a threshold 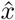 and increases to a higher value when 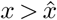. Only wealthy enough individuals (in terms of personal capital) can afford spending a large fraction of their resources into knowledge.

The consequences of this effect on cultural dynamics are shown in Figure 5 (**a-d**). Culture accumulates and has phases of acceleration (not shown), because the existing knowledge allows individuals to be culturally more productive. What is more, for reasonable parameter values, culture can be bi-stable. On the one hand, a population starting initially without any cultural knowledge accumulates little culture and eventually remains stuck in a “poverty trap” (Azariadis and Stachurski, 2005), at a very low level of knowledge. On the other hand, if the amount of cultural knowledge exceeds a threshold, for stochastic reasons for example or because of a change in the environment, then individuals reach a state where they can afford to invest a larger fraction of their resources in knowledge. They then manage to capitalize from generation to generation, which allows them to reach eventually a much higher level of cultural knowledge.

**Fig. 5.**
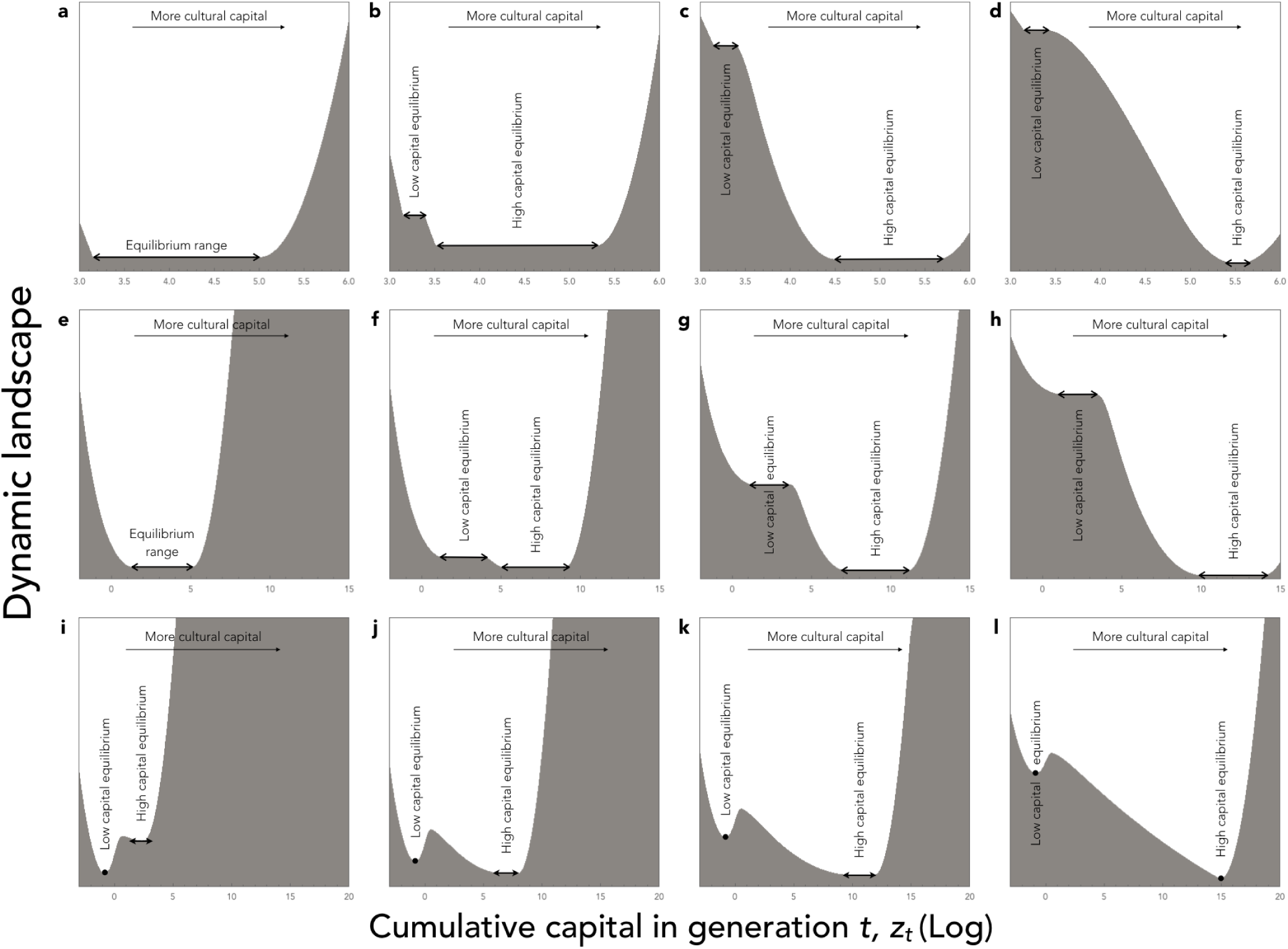
Cultural dynamics when culture is a capital. Graphic representation of cultural dynamics through an analogy with a physical landscape, as in figure 4. **a-d**: Investments in knowledge increase along the pyramid of needs, with parameters as in figure 3, but with *α* = (1 − *c*)*ζ* and *β* = *cζ*, where *ζ* = 0.9, and *c* is a hard threshold function of *x*: *c* = *c*_low_ for 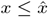, and *c* = *c*_high_ for 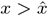, with 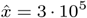, *c*_low_ = 0.05, and *c*_high_ = 0.05 (**a**), *c*_high_ = 0.1 (**b**), *c*_high_ = 0.5 (**c**), or *c*_high_ = 1 (**d**). **e-h**: Age at maturity increases with cultural capital, with parameters as in figure 3, but with the time available for growth, *L*, varying as a function of *z* according to a sigmoid function: 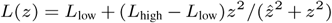, with 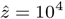, *L*_low_ = 10, and *L*_high_ = 10 (**e**), *L*_high_ = 30 (**f**), *L*_high_ = 50 (**g**), or *L*_high_ = 100 (**h**). **i** - **l**: With division of labor, and parameters as in figure 3, but with *α* = *β* = 0.5, and the division of labor model described in SI 2 with λ_min=0.01_, *ϕ* = *z, u* = 1, and the exponent of the production cost varying in function of *z* according to eq. 26 in SI, with *a*_high_ = 2, *a*_low_ = 1, and *s* = 10, and with population size *N* = 10 (**i**), *N* = 50 (**j**), *N* = 100 (**k**), or *N* = 1000 (**l**).

#### Parental care and age at first reproduction increase with the amount of cultural knowledge

The development of individuals depends, among other things, on two essential factors: the time during which they grow (that is, their age at maturity) and the amount of parental care they receive. Age at maturity is a life history decision of the offspring (Charnov, 1991, Stearns, 1977). Parental care is a life history decision of the parents who can choose either to produce a large number of offspring with few resources for everyone, or a small number of offspring with a lot of resources for each.

Empirical observations show that these two aspects of life history are plastic in humans, and depend in particular on the level of resources of individuals (Nettle, 2010, Roser, 2017). For instance, in the case of parental care, during their economic development, human societies make a relatively abrupt transition, the demographic transition, from a strategy with many children, each with relatively little investment, to a strategy with a high investment per child (Blanc and Wacziarg, 2019, Cummins, 2013). And a similar phenomenon occurs regarding the age at first reproduction. Economic development and increased levels of resources leads people, on average, to delay their reproduction and invest longer in their studies while, conversely, poverty leads women to choose to reproduce early (Roser, 2017).

We have introduced these two types of plasticity into our model. Formally, the age at sexual maturity is captured by the parameter *L*, and parental care is captured by our parameter *P*_0_ measuring the productivity of individuals that is independent of their own capital. In both cases, for simplicity, we assume that they increase with the level of resources of the parental generation, that is *L* and/or *P*_0_ in generation *t* + 1 increase with the amount of knowledge *z*_*t*_ available in the previous generation. Here we show results in the case where only *L* is plastic. We assume that *L*(·) is a sigmoid function, with a sudden change in strategy when *z*_*t*_ exceeds a threshold. This assumption leads, again, to bi-stable dynamics, with a sudden acceleration of cultural evolution when individuals’ life history strategy changes (Fig. 5 (**e-h**)). This occurs, here, because the knowledge of the previous generation directly accelerates the growth of the individuals of the next generation, and allows them to reach the frontier earlier in spite of the burden of knowledge.

We also included the possibility that the change of life history involved in the demographic transition may be accompanied by a reduction of population size. In reality, it has only been accompanied by a slowdown in population growth (not by a reduction of population size). But the important point is that there is indeed a trade-off between population size and parental care (or duration of growth) per child. Since population size has a small effect on the dynamics of cultural accumulation, however, this effect did not counteract the positive effect of life history (not shown). Cultural accumulation is much more intense in a small population where each individual has a lot of parental care and a long growth than in a large population where each individual has little care and little time to grow.

### Division of cognitive labor and the effective cost of learning

The efficiency of social learning itself can increase with the amount of cultural information available if, on average, the greater the amount of information produced in the past, the faster each unit of information is learned. This hypothesis is followed in many papers that assume the rate of social learning to be proportional to the amount of knowledge available that an individual does not yet not possess (e.g. Aoki et al., 2012, Lehmann et al., 2013). This assumption has no reason to be true in general. However, it is conceivable that sometimes it may be true, particularly if new cultural knowledge partially replaces old ones. Arabic numerals, for example, have replaced Roman numerals, and they are more useful but not more expensive to learn. The efficiency of social learning has therefore increased after the invention of Arabic numerals. More generally, people do not need to learn the whole history of science and technology, but simply to learn the latest state of the art. Besides, it is conceivable that part of cultural progress consists in improving existing knowledge by making it more compact, or easier to learn, and thus improving the efficiency of investments in social learning. Overall, therefore, the return on investment of social learning, λ, is likely to increase with *z*, which is a way for culture to increase the efficiency of individuals’ lives.

However, in fact, knowledge has a stronger and more general effect on the cost of learning through another mechanism. In practice, in any developed society with a large number of technologies, we acquire the benefits of the vast majority of cultural knowledge without having to learn it. For example, we benefit from our societies’ technical knowledge on internal combustion engines without having to learn anything, or almost anything, about the functioning, let alone the making, of these engines. Only a very small fraction of people in our societies invest in the acquisition of this knowledge. All the others obtain the benefits of internal combustion engines by exchanging with these specialists.

Thus, the true cost of “learning” a technique is not really the cost of learning it but rather the cost of acquiring the benefits of that technique via an exchange with others. This cost is lower because only one individual pays the cost of learning and then passes it on to all its customers. Standard models of cultural evolution take into account the existence of a division of labour of *innovation*, which constitutes the major benefit of culture in these models. But to capture the full benefits of culture, we must also take into account another division of labour: the division of labour of *social learning*.

This principle has an essential consequence. As technological progress creates economies of scale, it paradoxically reduces the effective cost of learning (that is, the cost of acquiring technical goods). The higher the level of technical knowledge in a society about food production, for example, the more a small number of producers can supply a large number of consumers at low prices. Thus, even if the cost of learning to become a farmer increases with the amount of technical knowledge (it requires longer and longer studies), the price of food can actually *decrease*, because this cost is passed on to a larger and larger number of consumers.

In SI 2, we formalize this principle with an approach inspired by micro-economic models of markets. The result of this analysis is illustrated in Figure 5 (**i-l**). In a large population, we observe once again bi-stable dynamics, with a poverty trap at low technical level (and low division of labour) and a sudden acceleration beyond a threshold, towards a new equilibrium at high technical level (and high division of labor). This occurs, here, because the knowledge of the previous generation reduces the effective cost of “social learning” for the next generation, which allows them to reach the frontier earlier in spite of the burden of knowledge. Besides, we also observe an indirect effect of population size as it constitutes an upper limit on the division of labour.

## Discussion

In this article, we have modeled cultural knowledge as a capital in which individuals invest at a cost. To this end, following other models of cultural evolution, we have explicitly considered the investments made by individuals in culture as life history decisions. Our objective was to understand what then determines the dynamics of cultural accumulation. Our models have shown that culture can accumulate provided it improves the efficiency of people’s lives in such a way as to increase their cultural productivity or, said differently, provided the knowledge created by previous generations improves the ability of subsequent generations to invest in new knowledge. And our central message has been that this positive feedback allowing cultural accumulation can occur for many different reasons, while the cultural evolution literature only considers one such reason. We have shown that it can occur if cultural knowledge increases people’s productivity, including in domains that have no connection with knowledge, because it simply frees up time that people can then spend learning and/or innovating (Figs. 3 and 5 (**a-d**)). We have shown that it can occur if cultural knowledge, and thus the higher level of resources that results from increased productivity, leads individuals to modify their life history decisions through phenotypic plasticity, either by delaying their sexual maturity or by producing fewer offspring but investing more in each (Fig. 5 (**e-h**)). And we have also shown that it can occur if technical knowledge reduces the effective cost of its own acquisition via division of labour (Fig. 5 (**i-l**)). In general, culture accumulates if it increases the *cultural* productivity of individuals, either by increasing their *overall* productivity, or by increasing the *fraction* of their production that they allocate to knowledge.

In the cultural evolution literature, culture can accumulate because it allows individuals to acquire knowledge and skills more efficiently than if they had had to create them from scratch (e.g. Aoki et al., 2012, Boyd and Richerson, 1995, Enquist et al., 2007, Enquist and Ghirlanda, 2007, Lehmann et al., 2013). Here, we show that the foundation of cumulative culture is actually broader than this. Culture can accumulate provided it makes life easier for individuals, giving them a head start and ultimately allowing them to go further than previous generations, and this can happen for many reasons, of which reducing the cost of acquiring knowledge is just one example. The added value of this generalized view of culture as capital is that it allows to account for a wider range of cultural dynamics. In particular, it can result in the accumulation of culture undergoing phases of acceleration, or being sometimes bistable. This is important because cultural evolution is known to have a wide diversity of dynamics, with periods of stagnation where the amount of knowledge is constant, periods of bursts where the amount of knowledge increases rapidly, and also periods of decline, where the amount of knowledge decreases (Aiyar et al., 2008, D’Errico and Stringer, 2011, Lehman, 1946, Mokyr, 1990; and see also Enquist et al., 2008, Kolodny et al., 2015 and references therein). So far, these dynamics have been explained in two ways: demography and interdependent discoveries.

1. The demographic explanation is based on the combination of two mutually reinforcing effects (see Aoki, 2015 for a review). (i) If there is more knowledge in a population, then individuals are more productive, which increases the carrying capacity of the environment and leads to a denser population. (ii) This increased population density, in turn, allows individuals to produce more new knowledge because “more heads means more ideas” (Aoki, 2015, Arrow, 1962, Henrich, 2004, Mesoudi, 2011). The combination of these two effects leads to a positive feedback loop that can cause an acceleration of cultural accumulation, concomitant with a population densification. Conversely, phases of cultural decline are caused by the two opposing effects, also reinforcing each other.
2. The “interdependent discoveries” explanation, on the other hand, assumes that the phases of cultural acceleration are caused by key discoveries (e.g. calculus) that then facilitate the realization of a large number of lesser discoveries (Kolodny et al., 2015, 2016). Cultural evolution thus takes place in a punctuated manner, with phases of stagnation, and sudden bursts when a key discovery arises. In principle, the same effect can also explain cultural regressions. If some key knowledge is lost leading to the disappearance of a large number of more applied knowledge.

Both of these explanations have problems, however.

1. The demographic explanation entails that the times and places where cultural evolution should be the fastest are the most densely populated and/or the places where local sub-populations are the most interconnected. Although not definitively invalidated, this prediction is not readily in line with observations. Archaeological and ethnographic observations are often inconclusive or even sometimes at odds with the demographic explanation (see Aoki, 2015 and Vaesen et al., 2016 for reviews). And historical data (Allen, 2009, Clark, 2008, Mokyr, 2016) show that small populations (e.g. the UK) often created more innovations than larger ones (e.g. medieval China) and that, in a given country, periods of cultural acceleration do not coincide with significant demographic rises (e.g. China during the 19th century).
2. The “interdependent discoveries” explanation predicts that cultural accelerations, or decline, should be domain-specific. For instance, knowledge in physics should accumulate rapidly after the discovery of calculus but this should have no impact on biology or paintings. However, observations show that, on the contrary, the historical periods and areas of the world that were flourishing were flourishing in all domains, or in a large number of domains at the same time, including in completely unrelated areas (physics, biology, gastronomy, paintings, music, etc.). Classical Greece, Abbasid Irak, Song China or Renaissance Italy naturally come to mind.

Several scholars have already put forward the general importance of creativity in determining the pace of cultural evolution (Fogarty et al., 2015, and see also Kobayashi and Aoki, 2012), and others have even suggested that cultural acceleration could result from the fact that, for a reason or another, cultural knowledge itself increases either creativity (Enquist et al., 2008, Mesoudi, 2011) or the efficiency of social learning (Mesoudi, 2011). But these scholars did not propose a specific reason why this last effect should occur. In this paper, we propose a straightforward explanation of this effect. We propose that cultural knowledge simply creates resources, and that resources incites individuals to invest more into knowledge, generating a positive feedback. We show that this simple mechanism explains that the rate of cultural evolution (i) is largely independently of demography, and (ii) changes through historical time in independent areas of knowledge in parallel.

With this model, and more generally with all models in which cultural evolution depends on the resource allocation decisions of individuals (Aoki et al., 2012, Kobayashi et al., 2019, Lehmann et al., 2013, Mullon and Lehmann, 2017, Ohtsuki et al., 2017, Wakano and Miura, 2014), the field of cultural evolution comes close to a branch of economics, called endogenous growth theory, that also seeks to understand the conditions allowing the long-term accumulation of knowledge (Arrow, 1962, Jones and Manuelli, 1997, Romer, 1994, 1986). An important distinction between the two approaches, however, lies in the way they treat the life history of individuals. In models in endogenous growth theory, or at least in the foundational models of this field, the life history of individuals –that is, firms and consumers– is considered an exogenous factor, whereas evolutionary biologists typically aim to consider cultural accumulation as the consequence of investment decisions that are part of the life history strategy of individuals. This explicit consideration of life history allows us to highlight the fact that biological reproduction is a waste from the point of view of cultural evolution, as newborn individuals have to reach the frontier from scratch, which constrains significantly the possibility of accumulating knowledge. A similar waste is taking place in endogenous growth theory in the form of consumption –as opposed to investment– and plays a key role in the accumulation of knowledge. In a way, then, one way to see the merit of evolutionary approaches is that they provide a biological foundation for the trade-off between consumption and investment that economists consider key to economic progress.

Our article has a number of methodological limitations that need to be acknowledged. The first concerns the extent of phenotypic plasticity. We have assumed that individuals are always optimal in their decisions about resource allocation. This implies that they are endowed with *perfect* phenotypic plasticity. In reality, especially in environments that have been profoundly modified by culture, individuals probably make mistakes in their allocation decisions, under-investing for example in innovations (or the other way around) and it would be important to characterize these mistakes and take them into account in further models.

Second, it is worth noting that our model primarily deals with technical or scientific types of knowledge that have a positive effect on the vital rates of individuals (including on their ability to produce subsequent knowledge). Its generalization to “symbolic” cultural phenomena such as art, narratives, religious representations, games, sports, etc. is warranted only to the extent that they confer immediate benefits for individuals.

Third, and most importantly, our model is deterministic. We do not consider the stochastic aspects of cultural evolution. This limitation has three dimensions. First, we consider innovations and learning as deterministic growth processes when in reality they have a stochastic component. Second, we consider a mean-field model in which all individuals are identical. In reality, individuals always differ at least slightly from each other, so that exceptionally creative individuals may appear sporadically in populations. Finally, we model culture as a single quantitative dimension, whereas a more realistic model should explicitly consider different cultural items, which would allow capturing the fact that key discoveries can suddenly accelerate the emergence of other discoveries (Kolodny et al., 2015, 2016). These three types of stochastic mechanisms are likely to cause shocks in cultural evolution. They could play an important role in the dynamics of culture and explain some of the variability in the trajectories of different populations.

To conclude, a potentially interesting aspect of our approach is the way it makes us think about culture. If culture can accumulate because, very generally, it improves people’s lives, then it is not an area of production in isolation from other human production. Culture is a capital that is transferable to subsequent generations in the same way that other forms of capital can be transferred as well. Hence, we began this article by arguing that cultural knowledge should be considered as part of the *heritable* capital produced by individuals, this form of capital differing from *non-heritable* capital in that it retains its value after the death of its producer. But we would like to suggest now that it could be useful in fact to broaden the definition of culture to include in fact *all* forms of heritable capital, including those that are not knowledge. The knowledge and skills of a population (i) can accumulate, (ii) have lasting effects over time and on a large number of individuals and (iii) partly explain the differences between human societies. But other properties of the environment, also resulting from the efforts of past human beings, have exactly the same effects. An area that has been deforested around a village, a temple or church built by ancestors, a reduced density of dangerous predators, an efficient railway system, a trustworthy political institution, etc.; all these human productions have a lasting influence on people’s development and behaviour, can accumulate, and explain a part of human variability, just as knowledge does. Hence, the culture of a society, we think, has no reason to be restricted to knowledge and skills but should be seen, more broadly, as a capital produced by past generations that affects the lives of present generations.

## Acknowledgments

We thank Laurent Lehmann and two anonymous reviewers for very helpful comments on a previous version of this manuscript. This work is supported by ANR-17-EURE-0017 FrontCog and by ANR-10-IDEX-0001-02 PSL.

## Supporting Information

### 1. Evolutionarily stable schedule

#### 1.1. Model 1

In model 1, the production function is merely a target that individuals seek to maximize when they invest in different compartments of their capital, but this model deliberately neglects the fact that this function in turn determines the amount of resources that can be invested per unit of time in different forms of capital. It is assumed that, whatever the amount of capital owned by an individual, she produces the same constant amount of resource per unit of time (without loss of generality, this constant is considered equal to 1). This is a deliberately false approximation but it serves as a point of comparison with the next model.

In both models, we assume that individuals follow a bang-bang life history strategy, investing in their growth for a duration *L* and then in reproduction for the remaining of their life –the duration of which needs not be specified in the model. In principle the time *L* that individuals devote to growth, before they start to reproduce, should itself be a decision of individuals, evolving by natural selection. However, in the sake of simplicity, we neglect this aspect of resource allocation in both models, and assume that individuals have an exogenously fixed time, *L*, available to grow. In this case, the evolutionarily stable strategy simply consists in aiming to maximize one’s productivity at age *L*, as this fully determines one’s total reproductive success. As long as the production function is concave (which is the case here since we have assumed *α* ≤ 1 and *β* ≤ 1), standard results of optimal control theory (see Iwasa and Roughgarden, 1984, Perrin, 1992, Perrin et al., 1993) tell us that the optimal allocation strategy for individuals to do so consists in investing at all times in the activity (growth, learning, or innovation) that has the greatest marginal impact on productivity (see Iwasa and Rough-garden, 1984, Perrin, 1992, Perrin et al., 1993). From the production function given by the equation 1 of main text, the marginal production of *x* and *y*, respectively, are given by

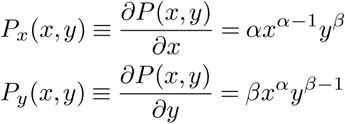

So the marginal return of investing a unit of time in growth is *γP*_*x*_(*x, y*), the marginal return of investing a unit of time in learning is λ*P*_*y*_(*x, y*) and the marginal return of investing a unit of time in innovation is *ηP*_*y*_(*x, y*) with *η* < λ.

We consider a population of *N* individuals developing in parallel in an environment in which an amount *z* of knowledge is already available from the previous generation.

##### 1.1.1. Phase 1: Learning and growth

As long as individuals have not learned all the cultural information already available (that is, as long as *y* < *z*_0_) then the return on innovation is always lower than the return on social learning, and individuals thus only arbitrate between (i) growth and (ii) social learning. During this phase, if *γP*_*x*_(*x, y*) > λ*P*_*y*_(*x, y*) then individuals prefer to invest all their resources in growth, while if *γP*_*x*_(*x, y*) < λ*P*_*y*_(*x, y*) then they prefer to invest all their resources in social learning. Natural selection therefore favours a strategy that consists of investing a fraction of the resources in growth and a fraction in learning in order to maintain at all times *γP*_*x*_(*x, y*) = λ*P*_*y*_(*x, y*), which implies maintaining a ratio

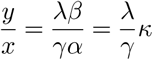

where *κ* ≡ *β/α*. To maintain this ratio, individuals invest a fraction of their resources *l* in social learning and a fraction 1 − *l* in growth, with

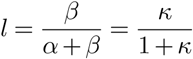

The growth rate of *x* during this phase is therefore *γ*(1 − *l*) while the growth rate of *y* is λ*l*. This phase lasts until all available knowledge has been learned, so until *y* = *z*, which takes a time

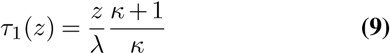

At the end of this phase, we have *y* = *z* and 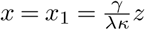.

##### 1.1.2. Phase 2: Pure growth

Once *y* = *z*, individuals cannot improve *y* through social learning. They only have the choice between innovating or growing. However, the efficiency of innovation is lower than the efficiency of social learning (i.e. *η* < λ), which implies *ηP*_*y*_(*x, y*) < λ*P*_*y*_(*x, y*). At the end of the first phase we had equal returns λ*P*_*y*_(*x*_1_, *z*) = *γP*_*x*_(*x*_1_, *z*). This implies *ηP*_*y*_(*x*_1_, *z*) < *γP*_*x*_(*x*_1_, *z*). That is, at the end of Phase 1, investing in growth is necessarily more profitable than innovating. So individuals do not start innovating right away. They first experience a phase during which they only invest in the growth of *x*. During this phase, the return of growth *γP*_*x*_(*x, y*) decreases. Hence, this phase ceases when this return meets that of innovation, that is when *γP*_*x*_(*x, y*) = *ηP*_*y*_(*x, y*), which implies 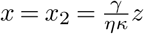, at which point individuals can start innovating (phase 3). This second catch-up phase therefore lasts for a duration

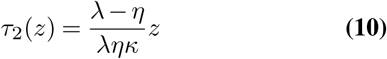

Hence, catching-up takes a total time *τ* (*z*) ≡ *τ*_1_(*z*) + *τ*_2_(*z*), which gives

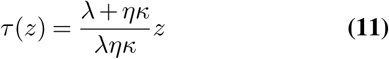

This allows us to define the average growth rate of individuals during the catch-up phase

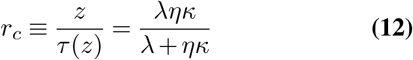

##### 1.1.3. Phase 3: Innovation, learning and growth, at the frontier

At the end of phase 2, the marginal return on investment in growth is exactly equal to the return on investment in innovation *γP*_*x*_(*x, y*) = *ηP*_*y*_(*x, y*). Since social learning is always more efficient than innovating (λ > *η*) this means that social learning is still the most profitable activity. So, as soon as an individual in the population produces an innovation, then all the other individuals prefer to invest in learning this innovation rather than in any other activity. In other words, during this third phase, individuals do not only invest in innovation and growth, they always keep a fraction (sometimes very large in fact) of their resources to invest in the social learning of *all* the innovations produced by others. The difficulty then consists in calculating the rate at which a population of *N* individuals produces new knowledge. To do this, let us assume that each individual invests a fraction of his resources *d* in innovation, a fraction *l* in social learning, and a fraction 1 − *l* − *d* in the growth of his personal capital *x*. Two conditions must be met by *d* and *l*.

1. Individuals must learn all innovations produced by others, which implies that *l* and *d* must follow the condition λ*l* = *ηd*(*N* − 1).
2. Individuals must maintain the respective returns of innovation and growth equal, i.e. *ηP*_*y*_(*x, y*) = *γP*_*x*_(*x, y*). This implies that they maintain the ratio 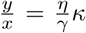. Therefore, *l* and *d* must also follow the condition 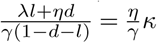.

These two conditions can be solved. This gives us 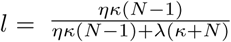 and 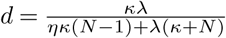. The speed of cultural evolution during this third phase is *r*_*i*_ ≡ λ*l* + *ηd* = *ηdN*, which therefore gives

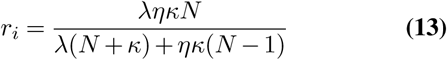

If a population contains an amount of knowledge *z*_*t*_ in generation *t*, in the next generation it contains an amount 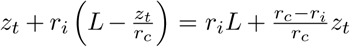. Considering the initial condition *z*_0_ = 0, this recurrence equation leads to the explicit dynamics of *z*_*t*_ over successive generations:

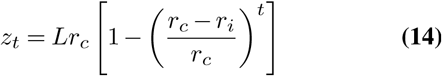

#### 1.2. Model 2

In model 2, it is explicitly considered that the production function determines the amount of resources available per unit of time, that individuals can then allocate to their different activities. Here as well, we assume that individuals have an exogenously fixed time, *L*, available to grow such that the evolutionarily stable strategy simply consists in aiming to maximize one’s productivity at age *L*. Here as well, as long as the production function is concave, the optimal allocation strategy for individuals is to make, at each instant, the investment(s) with the strongest marginal effect on productivity. This simple principle allows us to simulate the development of individuals.

The problem is that, in model 2, the growth rate of individuals is not constant (unlike model 1) but given by the production function and the development of individuals is therefore not analytically solvable. We therefore simulate numerically step by step this development in a social environment in which they initially have an amount *z* of available knowledge from the previous generation. These simulations are performed by choosing a time step *δ*, the smallest possible (ideally infinitely small). At each time step we numerically calculate the marginal return, *P*_*x*_(*x, y*) and *P*_*y*_(*x, y*), of each type of capital and we calculate the growth of individuals during this time step as follows:

1. When *y* < *z* (that is, individuals still have things to learn socially). (i) If λ*P*_*y*_(*x, y*) < *γP*_*x*_(*x, y*) individuals invest all their production in growth and therefore *x* increases by *γP* (*x, y*)*δ* while *y* remains constant. (ii) If λ*P*_*y*_(*x, y*) > *γP*_*x*_(*x, y*) individuals invest all their production in social learning and therefore *y* increases by λ*P* (*x, y*)*δ* while *x* remains constant.
2. When *y* ≥ *z* (that is, individuals no longer have anything to learn socially from the previous generation). (i) If *ηP*_*y*_(*x, y*) < *γP*_*x*_(*x, y*) individuals invest all their production in growth and therefore *x* increases by *γP* (*x, y*)*δ* while *y* remains constant. (ii) If *ηP*_*y*_(*x, y*) > *γP*_*x*_(*x, y*) individuals invest all their production in both (i) innovating and (ii) learning the innovations made by others. In this case, *y* increases by 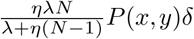 while *x* remains constant.

With these simulations, by considering a population of *N* individuals with a quantity of knowledge *z*_*t*_ available in generation *t*, we can simulate the development of individuals and thereby obtain the quantity of knowledge that they reach at the end of their development, which gives us *z*_*t*+1_, the quantity of knowledge available for the next generation. Eventually, performing the same simulations for different values of *z*_*t*_ allows us to plot *z*_*t*+1_ in function of *z*_*t*_ (Figs. 3 and 5 of main text).

In the particular case where the relative importance of *x* and *y* depends on the capital of individuals (section of main text), we assume that the two exponents, *α* and *β*, of the production function (Eq. 1 of main text) depend on non-heritable capital, via two functions *α*(*x*) = (1 − *c*(*x*))*ζ* and *β*(*x*) = *c*(*x*)*ζ*, with *ζ* ∈ [0.1] and *c*(·) an increasing monotonic mapping from [0. + ∞] to [0.1]. The model is then exactly the same except that the marginal productivities of *x* and *y* are given by

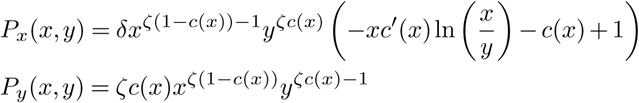

### 2. Division of labour

Here we consider a model in which individuals share the work of learning and producing cultural goods. We assume that they do so in a cooperative manner by seeking the most efficient division of labour possible. When the size of the population and the discrete nature of individuals does not constrain the division of labour, then the result under this assumption is identical to the long-term equilibrium of a competitive market (see Mas-Colell et al., 1995, chapter 10).

We consider a population of *N* individuals and we consider to simplify a good for which the demand is constant. The good has a benefit *b* and each of the *N* individuals in the population needs exactly one and only one token of this good. Among the *N* individuals in the population, *n* are producers of the good (and also consumers like everyone else). The ratio *N/n* therefore measures the extent of the division of labour. The higher it is, the more extensive is division of labour.

The cost of producing the goods is two fold. First, being a producer entails a “fixed cost”, *ϕ*, that is independent of the amount of goods actually produced, and represents the cost of acquiring the knowledge and skills to make the good, and more generally all the fixed costs necessary to become a producer. Second, being a producer entails a “production cost”, *π*(*q*), that depends on the number *q* of goods produced. Typically, the production cost function should be convex, at least at some point, such that it becomes too costly to produce more goods (see Mas-Colell et al., 1995, chapter 10). Here, we consider a standard production cost

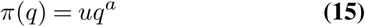

where *u* is the cost of producing one unit of the good and *a* > 1 measures the convexity of the production cost.

Let us first assume that there are exactly *n* producers of the good in the population. Every producer will have to produce exactly *N/n* goods that they sell at a price *p*. In this case, non-producing consumers earn a payoff *b* − *p*. And producers earn a payoff *g*(*p, n*) given by

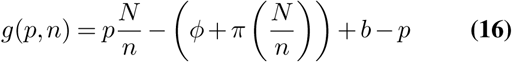

We assume that the number of producers, *n*, and the price of the good *p* are (i) in equilibrium and (ii) efficient. They must therefore comply with two conditions.

1. Equilibrium condition: The price of the good is such that producers and consumers earn the same payoff. Otherwise some consumers would prefer becoming producers, or vice versa (Mas-Colell et al., 1995, chapter 10.F), and *n* would therefore not be in equilibrium. This implies the condition 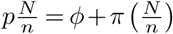, which gives us the equilibrium price 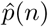 at which the good is sold when there are *n* producers:

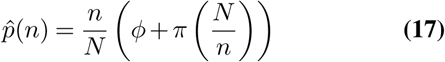 When the good is sold at the equilibrium price 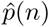, individuals share the cost of producing the goods in such a way that everyone earns the same payoff given by 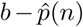.
2. Efficiency condition. The number of producers *n* should be such that the average payoff of individuals is maximized when the good is sold at the equilibrium price 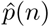. In combination with the condition 17 above, this implies that the 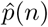 is minimized. So this gives us

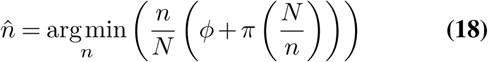

From these two conditions, and with the production cost given by equation 15, we obtain the optimal number of producers 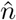 given by

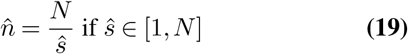

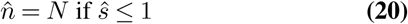

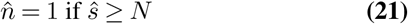

with

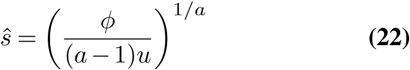

In other words, 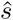 corresponds to the optimal division of labour in the absence of constraints (i.e. when the size of the population and the discrete nature of individuals are not limiting). 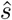 increases with the fixed cost *ϕ* and decreases with *a* and *u* which is expected since production techniques are less efficient. The actual division of labour is, however, constrained by (i) population size (at most all individuals will be producers) and above all by (ii) the discrete nature of the individuals (at least there must be 1 producer).

At the optimum division of labour, the price of the good is thus given by

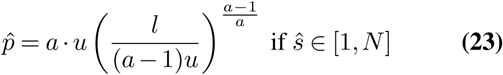

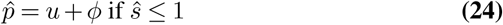

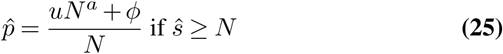

This prize is then integrated into the cultural evolution model as follows.

First, the fixed cost of becoming a producer, *ϕ*, represents the amount of cultural information needed to become a producer. We assume that this cost is simply given by *ϕ* = *z*. Second, we assume that the greater the amount of information, *z*, available in the previous generation, the more efficient are the techniques for producing goods. This is modelled by assuming that the convexity of the production cost function (eq. 15) is a decreasing function of the technical complexity *z*, equal to *a*_high_ when *z* = 0, and asymptotically reaching *a*_low_ when *z* tends toward infinity:

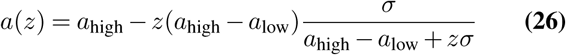

During the catch-up phase, when individuals still have less knowledge than the amount of knowledge available in the previous generation (*y* < *z*), the effective learning rate is given by 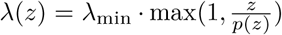. At worst, when the price of cultural goods is very high, the learning rate is λ_min_. When *z* increases, the efficiency of production techniques improves so that the division of labour is greater (the number of producers decreases), the price of goods decreases (*p*(*z*) < *z*), and thus the effective learning rate increases (i.e. λ(*z*) > λ _min_). When individuals are innovating at the frontier, however, then we assume that their learning rate is constant, and given by λ_min_.

